# Structure of a reversible amyloid fibril formed by the CPEB3 prion-like domain reveals a core sequence involved in translational regulation

**DOI:** 10.1101/2022.12.07.519389

**Authors:** Maria D. Flores, Michael R. Sawaya, David R. Boyer, Samantha Zink, Susanna Tovmasyan, Adrian Saucedo, Chih-Te Zee, Jorge Cardenas, Luana Fioriti, Jose A. Rodriguez

## Abstract

The cytoplasmic polyadenylation element-binding protein 3 (CPEB3) is a prion-like RNA-binding polypeptide. As a functional prion, CPEB3 is thought to modulate protein synthesis at synapses and enable consolidation of long-term memory in neurons. Here, we report that the prion-like domain 1 of CPEB3 self-assembles into labile amyloid fibrils *in vitro*. A cryoEM structure of these fibrils reveals an ordered 48-residue core, spanning L103 to F151. CPEB3 constructs lacking this amyloidogenic segment form abnormal puncta in cells when compared to wild type CPEB3, with reduced localization in dormant p-bodies and increased localization in stress granules. Removal of the amyloid core segment in CPEB3 also abolishes its ability to regulate protein synthesis in neurons. Collectively, this evidence suggests that the newly identified amyloidogenic segment within the CPEB3 prion domain is important for its regulated aggregation in cells and suggest its involvement in regulating translational activity and potentially long-term memory formation.

## Introduction

Prions, or proteinaceous infectious particles^1^, can form stable, long-lived pathogenic or functional assemblies in species throughout the tree of life^2^. Prion proteins are thought to rely on an amyloid architecture to recruit soluble proteins and template their conversion of into ordered fibrillar β-sheet-rich assemblies ^2,3^. Because of their ability to form amyloids, prion-like proteins present a challenge to the proteostatic framework in neurons^4^. However, since prions can endure protein turnover, they also represent a possible substrate for memory consolidation^5^. In fact, functional prions have been proposed to enable the persistence of memory engrams over time^6,7^.

A range of studies have provided evidence for prion-based memory consolidation mechanisms operating in neurons in *Aplysia*^8^, *Drosophila*^9,10^ and most recently, mammals^7,11^. Neuronal isoforms of the cytoplasmic polyadenylation element binding protein (CPEB) are involved in regulating localized translation at synapses^4^. Of the mammalian CPEB orthologs, CPEB3 distinctly contains a Q-rich N-terminal domain^10^, resembling those found in functional yeast prions^2^. CPEB3 is a 716-residue protein with two N-terminal prion-like domains linked by a short, low-complexity regulatory motif (Figure 1a). C-terminal to the CPEB3 prion domains are two RNA-binding domains and a zinc finger domain^10^ (Figure 1a). As a functional RNA binding protein, CPEB3 regulates translation of target mRNAs that are required for synaptic plasticity and the growth of new synaptic spines^11–13^. The prion-like nature of CPEB3 is supported by work showcasing its ability to form heritable aggregates in yeast^7^. Similar aggregates appear to be required for its function in neurons^7^. The N-terminal, prion-like domain 1 (PRD1) of CPEB3 is important to these traits as it rescues aggregation phenotypes in cells and mediates long-term potentiation (LTP) as well as the persistence of memory in mice^11^.

**Figure 1.**
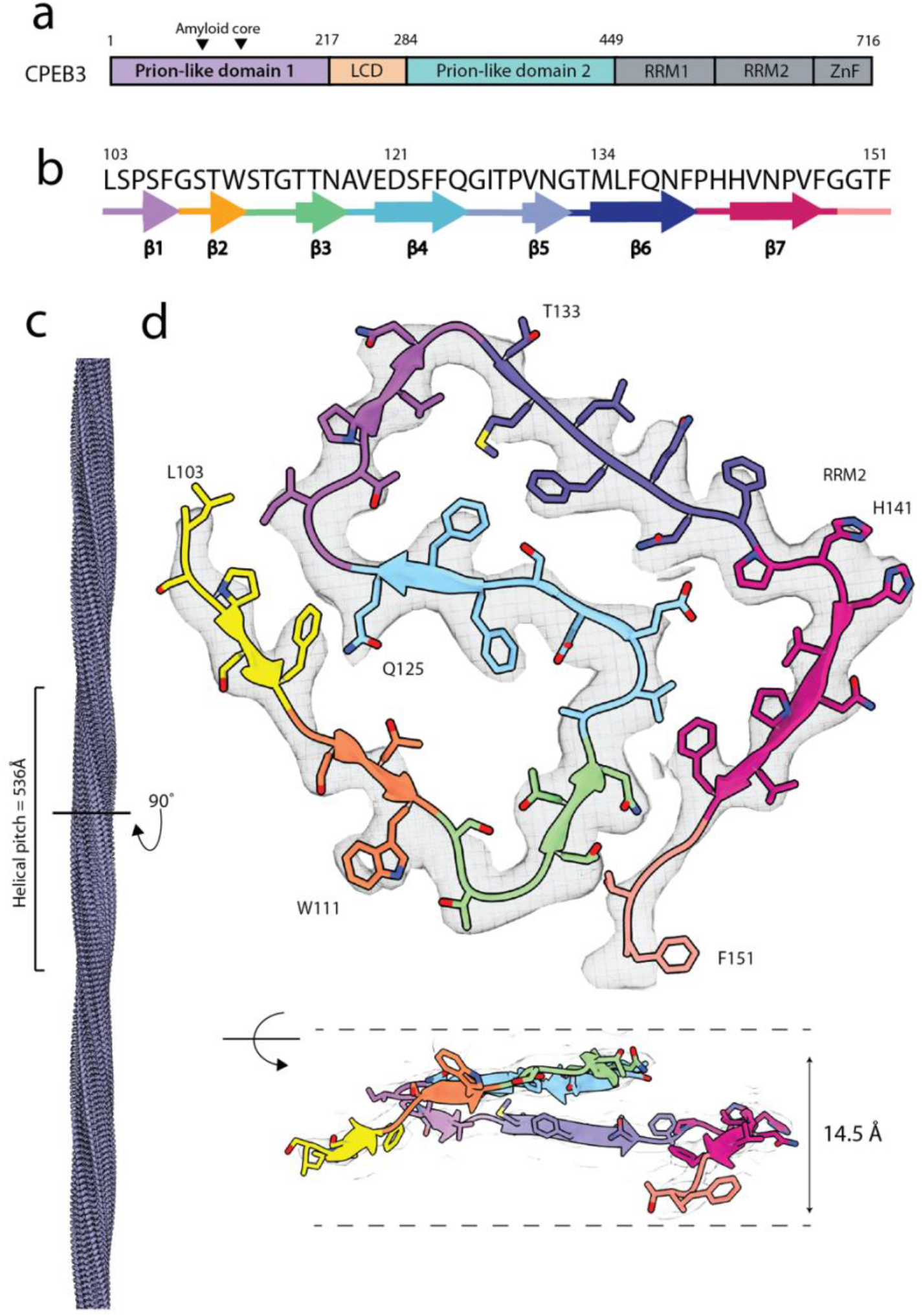
CryoEM structure of CPEB3’s prion-like domain. **a**, Schematic summary of CPEB3 domains. **b**, Amino acid sequence alignment of fibril core corresponding to secondary structure β-strands. **c**, Model of CPEB3AC filaments with a corresponding helical pitch of 536 Å. **d**, Top: cryoEM map of cross section perpendicular to the helical axis with model rendering colored to match secondary structure schematic. Bottom: side view of map and model measuring height different within one helical layer.

CPEB3 aggregates are tightly regulated and rely on mono-ubiquitination^12^ and SUMOylation^14^, which allow for proper activation and repression of the prion-like protein, respectively. In its basal state, CPEB3 is SUMOylated and colocalizes to p-bodies where it inhibits translation of target mRNAs^13^. Upon proper signal induced activity it is then shuttled to polysomes where it increases translation of its targets^13^. These observations suggest that CPEB3’s localization is directly related to its function.

A structure of fibrillar aggregates formed by the *Drosophila* CPEB ortholog, Orb2, isolated from fly brains, revealed a hydrophilic amyloid core with pronounced steric zipper motifs. The role played by this structure is consistent with the notion that CPEB can form persistent or stable amyloid assemblies, but how these assemblies form and how long they persist in cells remains unclear^15^. Additionally, while these studies present remarkable views of functional prion structures, they offer limited insights into the prion-like activity of mammalian CPEBs, and in particular CPEB3.^7–9^ Our current view of mammalian CPEB structure is instead informed by sequence analyses^16^ and NMR spectroscopy,^17^ which has revealed segments in PRD1 of CPEB3 that may form higher-order coiled-coiled α-helical assemblies. These efforts have contributed to the current delineation of certain CPEB domains as prion-like^11,7^.

We set out to understand the structural basis for prion-like activity by CPEB3 and found that recombinant CPEB3 PRD1 can self-assemble into reversible amyloid fibrils in solution. The structure of these fibrils, determined by single particle cryoEM, showed an ordered core composed of residues 103 to 151 of CPEB3 arranged in a serpentine fold. The sequence composition of this amyloid core (AC) offers clues to its lability and possible sensitivity to environmental changes in pH. CPEB3 constructs lacking this AC sequence showed altered subcellular distribution, indicating a role for this sequence in CPEB3 activity. This hypothesis is confirmed by the altered translational regulation of AC-deficient CPEB3 in primary neurons. These results suggest a model whereby CPEB3 function is regulated in part by the AC sequence in PRD1, possibly through its adoption of a labile amyloid fold.

## Results

### CPEB3 PRD1 forms reversible amyloid fibrils in vitro

Hypothesizing that CPEB3 PRD1 may form functional amyloid assemblies, we expressed and isolated a construct of human CPEB3 encoding its first 217 residues that included the entirety of its conserved PRD1 (Panel 1, Supplementary Figure 1). This region of CPEB3 has been previously identified as essential for prion-seeding, heritability^7^ and is essential for LTP in mice^11^. We assessed the ability of this construct (CPEB3_1-217_) to form fibrils *in vitro* by light scattering, enhanced fluorescence of the amyloid-binding dye ThT, and negative stain electron microscopy (Supplementary Figure 2). CPEB3_1-217_ tended to precipitate in denaturing solutions as amorphous aggregates, twisting fibrils, or flat, ribbon like sheets. In solutions containing 125 mM NaCl, 50 mM Tris-Base, 10mM K_2_HPO_4_ and 5 mM glutamic acid at pH between 4 and 5 (assembly buffer) CPEB3_1-217_ produced what appeared by negative stain electron microscopy to be fibrils with a homogenous twist (Supplementary Figure 2b). Solutions containing CPEB3_1-217_ fibrils exhibited enhanced ThT fluorescence (Supplementary Figure 2a). These fibrils were also labile, disassembling when diluted into water or exposed to low ionic strength solutions (Supplementary Figure 2c). To pursue structure determination in the face of this lability, fibrils had to be vitrified within an hour of formation in assembly buffer.

### Structure of the CPEB3 amyloid fibril core

Solutions containing CPEB3_1-217_ fibrils were interrogated by single particle cryoEM, with the goal of *de novo* structure determination (Supplementary Figure 4). CryoEM images confirmed that CPEB3_1-217_ forms rapidly twisting amyloid fibrils in assembly buffer, with a single polymorph representing over 90% of imaged particles (Supplementary Figure 5a). The remaining 10% of species exhibited variable twist or no twist, and therefore eluded further classification and structure determination (Supplementary Figure 5b). Two-dimensional class averages confirmed canonical amyloid features corresponding to the stacking of β-sheets separated by 4.8 Å interstrand distances, as well as a disordered flanking region, or fuzzy coat, surrounding the fibril core (Supplementary Figure 5b). Three dimensional reconstructions obtained from a subset of 40,329 segments yielded a 3.0 Å map that showed density for a single asymmetric protofilament that fit residues L103-F151 of CPEB3 (Figure 1b). We refer to this structure and its corresponding core sequence as the CPEB3 amyloid core (CPEB3_AC_).

CPEB3_AC_ fibrils carry a helical pitch of 536 Å and twist of -3.32° (Figure 1c). The length of the fibrils arises from stacking of CPEB3_AC_ molecules in parallel and in register intermolecular β-sheets. The tight serpentine fold adopted by the CPEB3_AC_ core is enabled by five sharp kinks interspersed among the molecule’s seven β-strands (Figure 1c). The fold exhibits two hydrophobic pockets and encloses three internal solvent channels. The CPEB3_AC_ core displays a total solvent accessible surface area per chain of 1653 Å^2^. At its N-terminus, L103 contacts a surface-facing segment of a beta turn in the fibril core, making a hydrophobic pocket composed of residues P105, F107, G126 and I127 (Supplementary Figure 7b). An additional hydrophobic pocket is formed by residues F124, V130, M134 and F136 (Supplementary Figure 7b). The core is further stabilized by stacking of aromatic residues, although some appear solvent facing, including W111, F139, and F151. Polar residues T110, S112 and T116 face the largest solvent channel buried within the fibril core. A second, smaller solvent channel is lined by a group of largely by hydrophobic residues: V119, P139, V143, P145; E120 stands as a lone hydrophilic residue in this group. This same glutamic acid forms part of a minute channel within the core, which is also lined by S122 and N138. The significance of E120 bridging these two solvent channels is unclear, but may indicate a role for it in the lability of the CPEB3_AC_ core.

### Structural features of labile CPEB_AC_ fibrils

Energy values calculated using coordinates of the CPEB3_AC_ fibril core show that, per layer, this core is less stable than those of other pathogenic amyloids (Supplementary Figure 7a)^18^. Estimates of its free energy are more in line with those of fibril structures formed by reversible, functional amyloids^19–21^. Solvent facing hydrophobic residues and buried solvent channels play a role in its predicted lability, but the CPEB3_AC_ is also strained by a series of sharp kinks. Some of these kinks tilt strands out of plane, warping the layers of the fibril to create a height difference of 14.5 Å across each of its rungs (Figure 1d, Supplementary Figure 7c). While the role of warping in amyloid fibril cores remains unclear, it may help relieve the strain associated with the large twist angle observed in CPEB3_AC_ fibrils. In addition, some kinks appear to be stabilized by local interactions. For example, a kink near G115 facilitates hydrogen-bonding between side chains of S112 and T116. Four of the 28 proline residues in CPEB3_1-217_ are resolved in the core. The remaining 24 proline residues are unresolved in the fuzzy coat of CPEB3_AC_ and may function to limit the size and stability of the ordered core. They appear as smeared density in 2D averages of CPEB3_AC_ fibrils (Supplementary Figure 4b). While some of these regions are hypothesized to form coiled-coils^16^, that region of the fibril remains insufficiently resolved to assign secondary structure.

### The role of negatively charged residues in CPEB_AC_

We observed four potentially charged residues in CPEB3_AC_: two histidine residues (H141 and H142) are solvent exposed, and two acidic residues (E120 and D121) are buried within the fibril core (Supplementary Figure 7a). The two histidine residues are located too far from the acid residues for charge complementation to stabilize the fold. E120 is located in the second solvent channel and is otherwise surrounded by hydrophobic residues, while D121 faces the opposing solvent channel (Supplementary Figure 7b). As CPEB3_AC_ aggregates near a pH of 5, E120 and D121, are anticipated to remain protonated within the fibril core and are capable of forming stabilizing hydrogen bonds. These same residues would render the core susceptible to pH triggered changes, as observed with other functional amyloids ^22,23^such as pmel17^22,23^ and β-endorphin^23^. To evaluate this hypothesis, we monitored the stability of CPEB3_AC_ fibrils in response to changing buffer conditions, using a ThT fluorescence assay. We found CPEB3_AC_ fibrils diminish in abundance as pH increases or as the ionic strength of the buffer decreases, denoted by a decrease of ThT signal and the absence of fibrils in negative stain images compared to fibrils that remain in their assembly buffer (Supplementary Figure 2c). This dissolution in response to changing environmental conditions is recapitulated by simulations tracking fibril stability over a period of 200 nanoseconds (Supplementary Figure 8a). Interlayer distances in a 5-layer fibrillar CPEB3_AC_ structure are disrupted as a pH increase from 3 to 7 is simulated; structural deviations are pronounced in charged residues E120 and D121 (Supplementary Figure 8b).

To assess whether E120 and D121 influence the morphology, distribution, and abundance of CPEB3 deposits in cells, we expressed a variety of GFP-tagged full-length CPEB3 constructs encoding wild type and E120/D121 variations, in HT22 and U87 neuronal cells (Figure 2a,b). In HT22 and U87 cells, wild type, GFP-fused CPEB3 formed fine, granular fluorescent puncta, as previously described (Figure 2c, Supplementary Figure 9a)^7,13^. Forty-eight hours after transfection, the fine puncta were replaced by larger puncta that were associated with reduced cellular health, likely due to prolonged CPEB3 overexpression (Supplementary Figure 9a). To probe the influence of acidic side chains at residues ^120^ED^121^ we expressed a CPEB3 variant encoding ^120^QN^121^, which retains the polarity and size of the wild type residues but ablates their negative charge. We also expressed a variant encoding ^120^AA^121^ to assess the effect of ablating polarity and reducing the size of wild type residues. Puncta formed by both constructs differed morphologically from those formed by wild type CPEB3-GFP under similar conditions (Figure 2c, Supplementary Figure 11). CPEB3-^120^QN^121^ generally formed irregularly sized and shaped puncta, while CPEB3-^120^AA^121^ formed fewer puncta that were intermediate in size and remained more regularly circular after 24hrs (Figure 2d, Supplementary Figure 10a, Supplementary Figure 11). The effects of these residue substitutions on the size and distribution of CPEB3^AC^ puncta implicate a functional role for E120 and D121.

**Figure 2.**
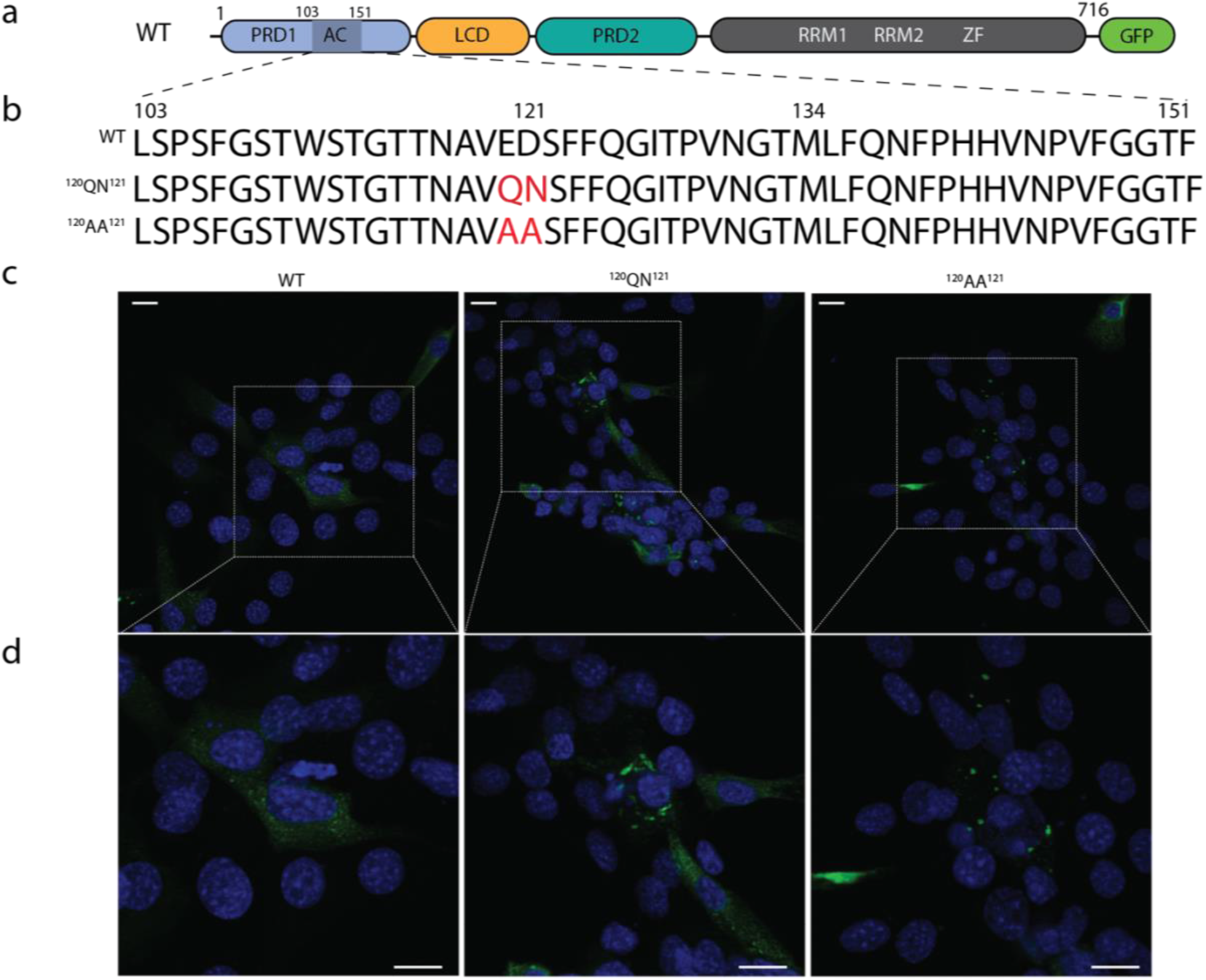
The role of E120 and D121 in CPEB3 puncta formation in cells. Wild type CPEB3-GFP shows fine, granular like aggregates in HT22 mouse hippocampal neurons after 24 hours of expression. **a**, Schematic of full-length WT construct expressed in cells. **b**, Sequence alignment of AC region in FL constructs with corresponding mutated residues in red. **c**, CPEB3-GFP WT and mutants (green) and nuclear staining (blue) in HT22 neurons. **d**, insets of panel c. Scale bar = 10 μM.

### CPEB3_AC_ is required for proper aggregation and localization in cells

To more generally assess the functional relevance of CPEB3_AC_ in cells, we asked whether the presence of the AC sequence affected CPEB3 aggregation. While CPEB3 is known to form aggregates of various sizes when overexpressed in cells^7,11,13^, a construct of CPEB3 lacking its amyloid core, CPEB3_ΔAC_ displayed a reduced capacity to form small aggregates (Supplementary Figure 9). Instead, this construct showed a delayed onset of larger aggregate formation as compared to wild type CPEB3 in both HT22 and U87 cells (Supplementary Figure 9). This prompted us to determine whether the localization of CPEB3_ΔAC_ differed from that of wild type CPEB3 in both cell lines. Under normal conditions CPEB3 is expected to localize in p-bodies^13^ that contain mRNAs and associated regulatory proteins. There, mRNAs are stored in a dormant state while they are trafficked to their destination to be translated, or can be degraded. Alternatively, CPEB3 could reside in Stress granules (SG), induced to promote cell survival by condensing translationally stalled mRNAs. We observed that wild type CPEB3-GFP aggregates primarily colocalized with the de-capping coactivator and p-body marker, Dcpa1, in unstimulated cells (Figure 3b). In contrast, in mouse HT22 hippocampal neurons, CPEB3_ΔAC_ aggregates primarily colocalized with the SG assembly endoribonuclease, G3BP, and minimally colocalized with the p-body marker, Dcpa1 (Figure 3b). Similar localization patterns were observed in human U87 glioblastoma cells (Figure 3c).

**Figure 3.**
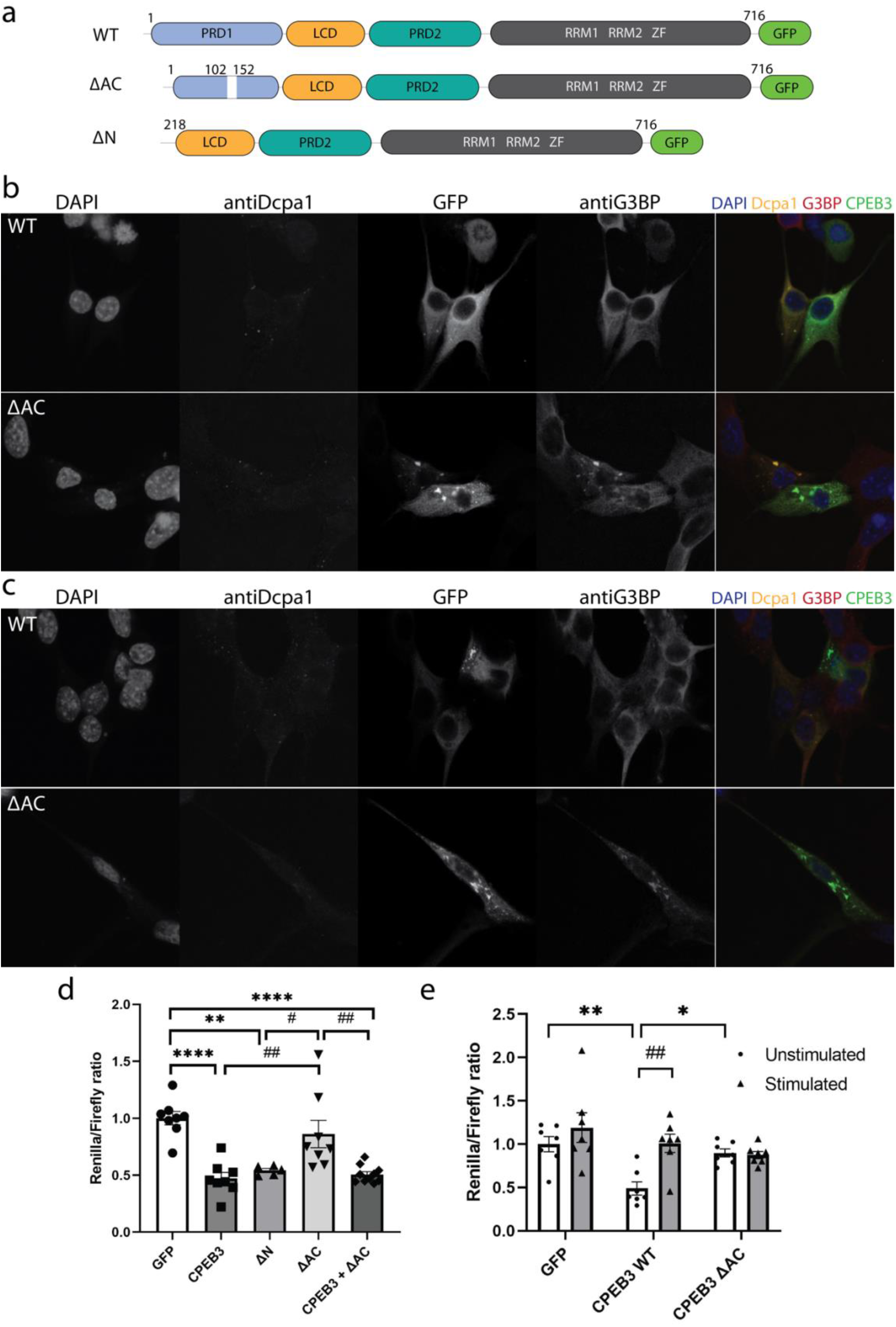
CPEB3_AC_ regulates subcellular location of GFP-tagged CPEB3 in cells and affects its translational regulation. **a**, Schematic representation of full-length constructs used in localization and translational assays. **b**, HT22 cells expressing WT (top) and AC (bottom) (green) and subsequent co-localization with antiDcpa1 (yellow) and antiG3BP (red) with nuclear stain (blue). **c**, Identical experiment to Figure 1b using U87 cells. **d**, HEK 293T cells inhibitory renilla luciferase assays (n =5-11 replicates per construct).) Graphs show the quantification of renilla/Firefly ratio, mean ± s.e.m. One-way ANOVA, followed by Tukey’s multiple comparisons post hoc test, ** p<0.01, ****p<0.0001 vs GFP; # p< 0.05, ## p<0.01, vs CPEB3^ΔAC^. **e**, Renilla luciferase assays performed in primary neurons. (n=7 per condition). Graphs show the quantification of renilla/Firefly ratio, mean ± s.e.m. Two-way ANOVA, followed by Tukey’s multiple comparisons post hoc test, * p< 0.05, ** p<0.01, vs CPEB3 unstimulated; ##p< 0.01 vs CPEB3 stimulated.

### CPEB3_AC_ impacts translational regulation by CPEB3

In its basal state, CPEB3 represses translation of target mRNAs, is SUMOylated, and colocalizes to p-bodies. After neuronal stimulation CPEB3 undergoes de-SUMOylation, leaves p-bodies and is then shuttled to polysomes to promotes translation of targets^13^. The increase in translational activity of CPEB3 is thought to be facilitated by monoubiquitination of PRD1 by Neuralized-1^12^. The ubiquitination site of CPEB3 is currently unknown and efforts to identify and characterize all SUMOylation sites^14^ have not yielded a complete understanding of their role in CPEB3 activity. Since CPEB3_AC_ is situated within PRD1, we postulated it may influence translational regulation by CPEB3. To test this hypothesis, we assessed translation of CPEB3 targets, comparing the impact of wild type protein to CPEB3_ΔAC_. Translation of Renilla luciferase fused to the 3’UTR of two established targets of CPEB3 targets, GluA2 and SUMO2 3’ UTR, in HEK-293 cells, was strongly inhibited by both wild type CPEB3 and CPEB3 lacking PRD1 (ΔN) (Figure 3d). In contrast, translation of the same targets was unaffected by CPEB3_ΔAC_; a marked change in translational regulation in the absence of the amyloid core segment. Co-expression of wild type and CPEB3_ΔAC_ in cells re-established the translational repression observed with the wild type protein alone, indicating CPEB3_ΔAC_ does not act in dominant negative fashion (Figure 3d).

Expression of wild type CPEB3 in primary neurons repressed translation of its targets in unstimulated cells (Figure 3e). To stimulate neurons, we used a protocol consisting of bath application of bicuculline and Glycine, a chemical treatment that induces long-term potentiation (LTP) of synapses (cLTP). As expected, glycine-mediated cLTP induced the translation of CPEB3 targets (Figure 3e). In contrast, expression of CPEB3_ΔAC_ under the same conditions ablated all translational regulation, rendering the protein essentially inactive in this capacity (Figure 3e). In fact, luciferase activity in the presence of CPEB3_ΔAC_ was most similar to that of cells expressing GFP alone, indicating that CPEB3_AC_ is required for regulated translation of CPEB3 targets.

## Discussion

CPEB3, a member of the cytoplasmic polyadenylation family of proteins, binds mRNA transcripts to activate or repress protein synthesis^24,25^. This process is hypothesized to be regulated by its prion-like self-assembly^4^, although the formation of prion-like structures by CPEB3 remains hypothetical. We find that a segment in the first CPEB3 prion-like domain forms labile amyloid fibrils. The ordered core of fibrils formed by this segment, CPEB3_AC_, is composed of a single asymmetric protofilament with parallel, in register strands. While the overall architecture of CPEB3_AC_ resembles that of a canonical amyloid structure, it contains buried polar residues and solvent channels that may decrease its stability. The ready assembly and disassembly of these fibrils would be a feature of CPEB3 assemblies in their role of translational regulation.

The amino terminal domains of many CPEB proteins are identified as prion-like in part due to their low-complexity sequences and their high number of uncharged polar residues^4,8,26^. Orb2 contains prion-like domains that form three-fold symmetric amyloid assemblies as isolated from fly brains^15^; these fibrils contain only a short 30-residue segment at their core. Likewise, in CPEB3, a short 48-residue segment within its first prion domain can form amyloid fibrils. However, some important features distinguish CPEB3 from Orb2 fibrils. First, while the PRD1 of CPEB3 encodes a low complexity sequence, CPEB3_AC_ is not particularly rich in Q or N residues. Also, unlike pathogenic amyloid fibrils, and perhaps unlike Orb2 fibrils, CPEB3^AC^ is readily labile. Features that may contribute to the lability of CPEB3_AC_ include its buried charged residues (E120, D121), its four prolines, and its highly warped layers. The buried charged residues would allow CPEB3_AC_ to respond to pH changes ^27^, enabling environmental queues to possibly regulate assembly of amyloid-like CPEB3_AC_ structures in neurons alongside other post-translational modifications involved in CPEB3 regulation in cells^12–14^.

The relevance of CPEB3_AC_ to the physiological role of CPEB3 is underscored by the fact that translational regulation by CPEB3 is ablated in cells that lack its amyloid core. The localization of CPEB3 to p-bodies is also sensitive to perturbations of CPEB3_AC_. In the absence of its amyloid core, CPEB3 localizes to stress granules, whose morphological appearance resembles those of other phase separated compartments^28^. The structural nature of CPEB3 within these aggregates, and whether it adopts an amyloid fold, remains unclear. CPEB3_AC_ may act as a regulatory element or may itself be subject to regulatory modifications such as SUMOylation^14^ and ubiquitination^12^. Such modifications may enable proper subcellular localization of CPEB3 and help avoid wanton self-assembly. This hypothesis would explain why ablation of this sequence causes loss of translational activity and the formation of large puncta. While the structures of amyloid fibrils reconstituted *in vitro* are known to be distinct from those isolated from disease tissue^29–33^, the relevance of CPEB3_AC_ to physiological CPEB3 function remains supported by: 1) the role of its AC sequence in localization to puncta in cells; 2) the fact that point mutations ablating charges in CPEB3_AC_ alter CPEB3 distribution in cells; and 3) that deletion of CPEB3_AC_ eliminates its ability to repress or activate mRNA targets, highlighting its role in translational activation by CPEB^7,11^ (Figure 4).

**Figure 4.**
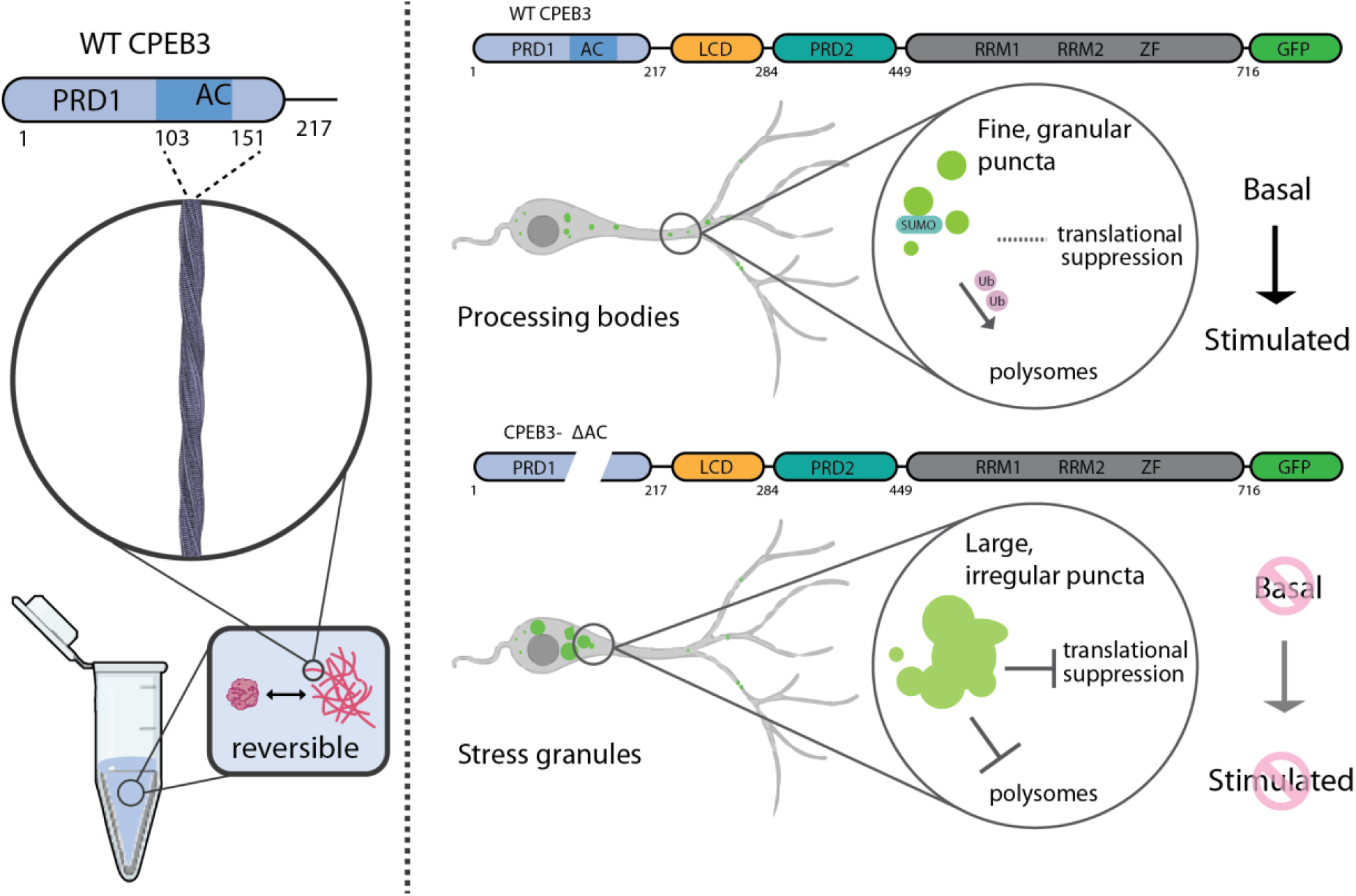
Model of CPEB3 AC and its role in CPEB3 aggregation and regulation. (Left panel) Amyloid fibrils formed in-vitro are reversible. Cryo-EM identifies a core sequence within the CPEB3 prion-like domain. (Right panel) WT CPEB3 displays fine, granular puncta that colocalize in p-bodies in the basal state and are moved to polysomes upon stimulation. CPEB3_ΔAC_ aggregates are larger and co-localize with SG markers, displaying no functional activity upon stimulation.

Our observation of CPEB3_AC_ provides additional experimental evidence for an expanding model of functional prions^4,34^ in which a labile amyloid state has evolved to perform biological functions^15,20,26^. Future insights into the prion-like role of CPEB3 in neurons and its regulated assembly and disassembly in cells will continue to expand our fundamental understanding of functional prion proteins. Ultimately, the structure of CPEB3 assemblies and their interacting partners in the brain remain important targets in the quest to understand the molecular basis for cellular memory.

**Table 1.** Cryo-EM data collection, refinement and validation statistics.

|  |  |
| --- | --- |
|  | CPEB3<sub>AC</sub> |
| **Data collection and processing** |  |
| Magnification | 105,000 |
| Voltage (kV) | 300 |
| Electron exposure (e-/Å<sup>2</sup>) | 52 |
| Defocus range (µm) | 1-3 |
| Pixel size (Å) | 0.86 |
| Symmetry imposed | C1 |
| Helical rise (Å) | 4.8 |
| Helical twist (°) | -3.32 |
| Initial particle images (no.) | 2,410,032 |
| Final particle images (no.) | 40,329 |
| Map resolution (Å) | 3.0 |
| FSC threshold | 0.143 |
| Map resolution range (Å) | 200-3.0 |
| **Refinement** |  |
| Initial model used (PDB code) | De novo |
| Model resolution (Å)<br>FSC threshold | 0.143 |
| Model resolution range (Å) | 200-3.0 |
| Map sharpening *B* factor (Å<sup>2</sup>) | - 100.35 |
| Model composition |  |
| Non-hydrogen atoms | 374 |
| Protein residues | 49 |
| Ligands | 0 |
| *B* factors (Å<sup>2</sup>) |  |
| Protein |  |
| R.m.s. deviations |  |
| Bond lengths (Å) | 0.006 |
| Bond angles (°) | 0.753 |
| **Validation** |  |
| MolProbity score | 1.79 |
| Clashscore | 9.26 |
| **Ramachandran plot** |  |
| Favored (%) | 95.74 |
| Allowed (%) | 4.26 |

---

**Supplemental Figure 1.**
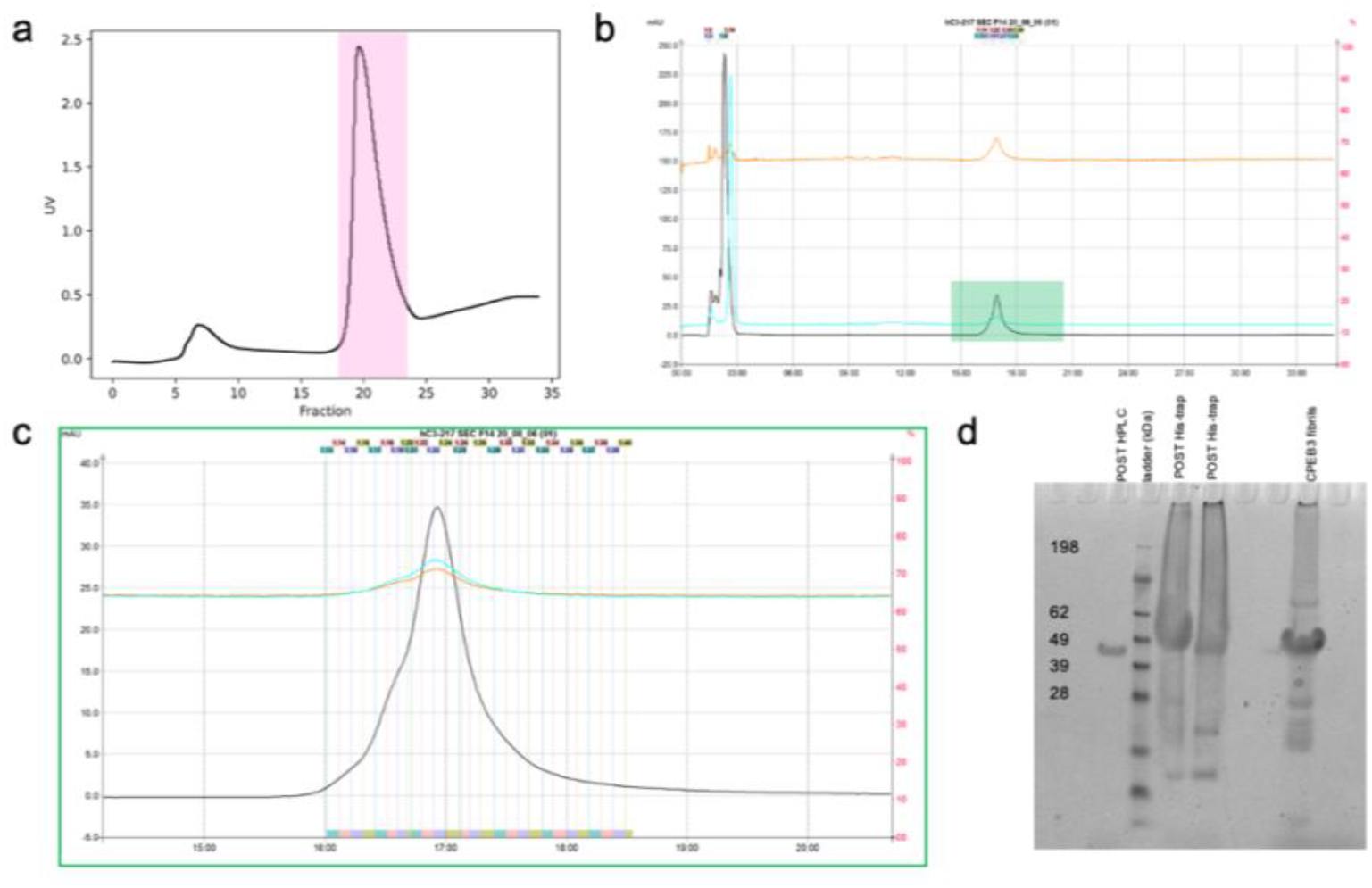
Recombinant protein purification.**a**, Example His-trap elution profile. CPEB3^1-217^ eluted in fractions 18-23 (pink subset). **b**, Full view of purification chromatogram. Three wavelengths were collected to monitor the purification: 214nm (black), 280 nm (orange), and 254 nm (light blue). **c**, Zoomed purification chromatogram (green) where the target protein started eluting. Three wavelengths were collected to monitor the purification: 214nm (black), 280 nm (orange), and 254 nm (light blue). **d**, SDS-PAGE shows harvested fractions after His-trap and HPLC purification. Freshly harvested fibrils also present.

**Supplemental Figure 2.**
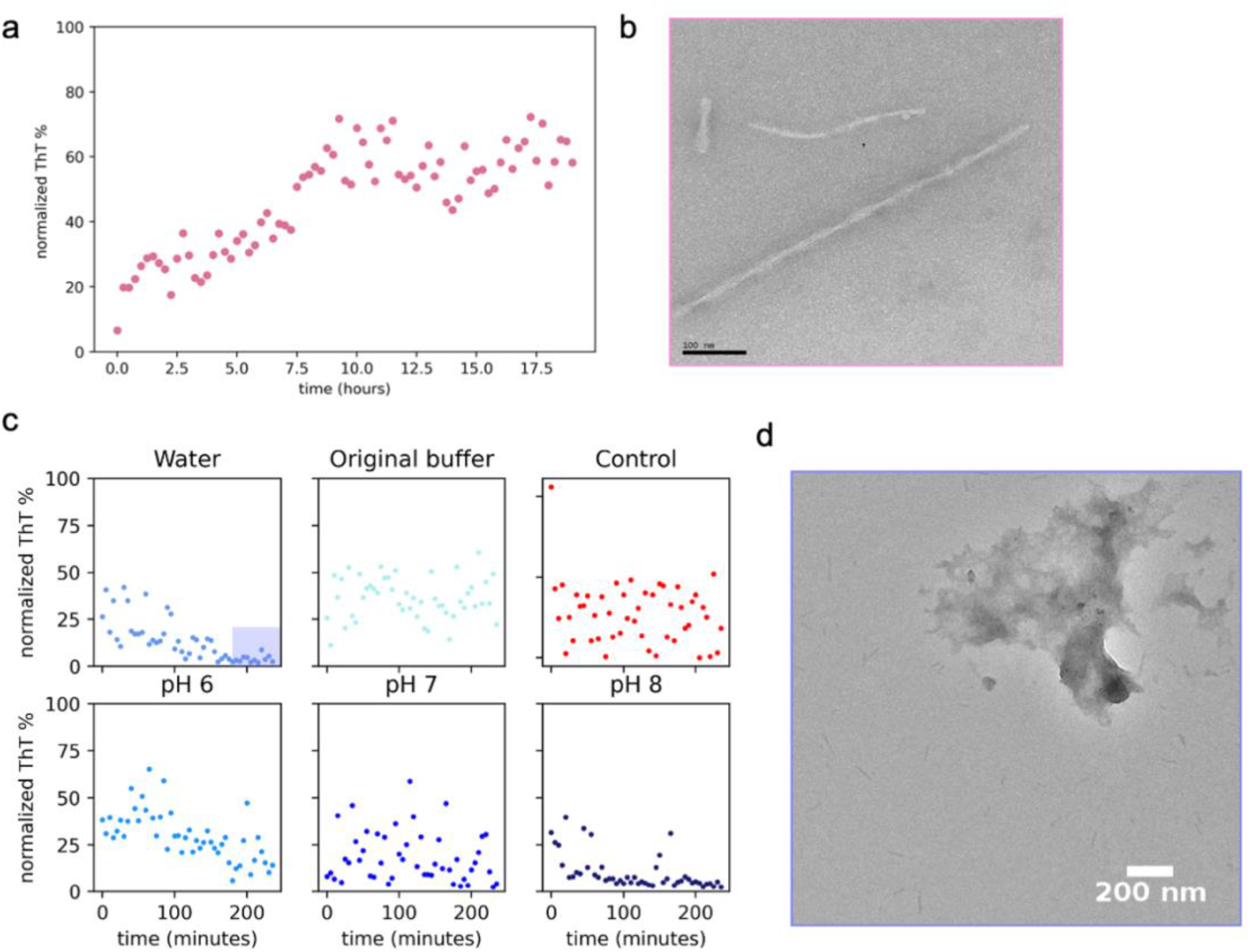
Biochemical characterization of CPEB3^1-217^ segment.**a**, ThT curves of CPEB3^1-217^. Data shown are normalized triplicate conditions, n=3. **b**, Negative stain TEM images of fibrils formed by CPEB3^1-217^. **c**, Dissolution assays of pre-formed fibrils. Samples were harvested after formation from SF 2a, and diluted 2-fold in corresponding buffers for approximately 4 hours. Data shown n= 4 wells per condition. **d**, Negative stain TEM image of fibril sample after dilution in water for 4 hours (purple subset in SF 1d).

**Supplemental Figure 3.**
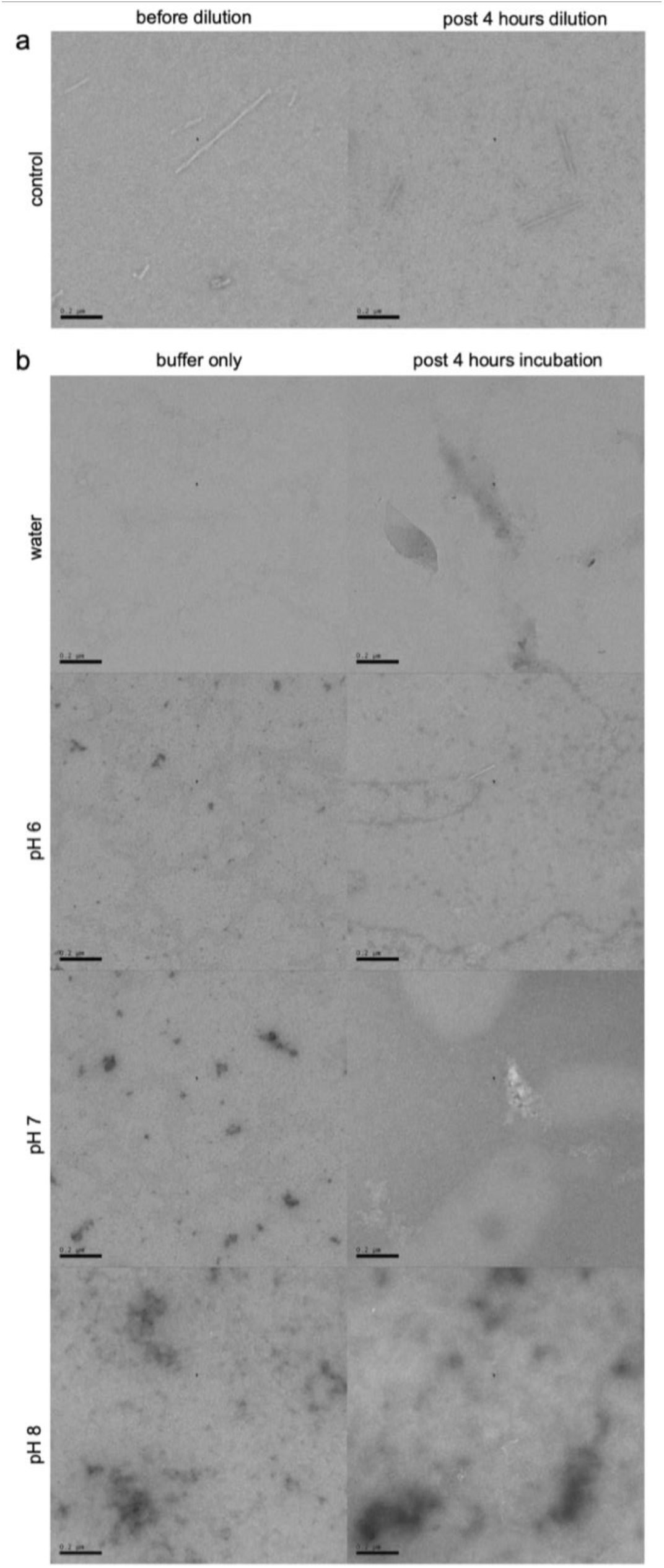
Negative stain TEM micrographs of CPEB3 fibrils after dissolution assays. Samples harvested from dissolution assays in Supplemental Figure 1d were prepped for TEM analysis.**a**, Control wells were diluted in original fibrilization buffer. **b**, Representative micrographs of blanks corresponding to blank, dilution buffer (left column), and protein samples after a 4-hour incubation in designated buffer (right column). Scale bar = 200 nm.

**Supplemental Figure 4.**
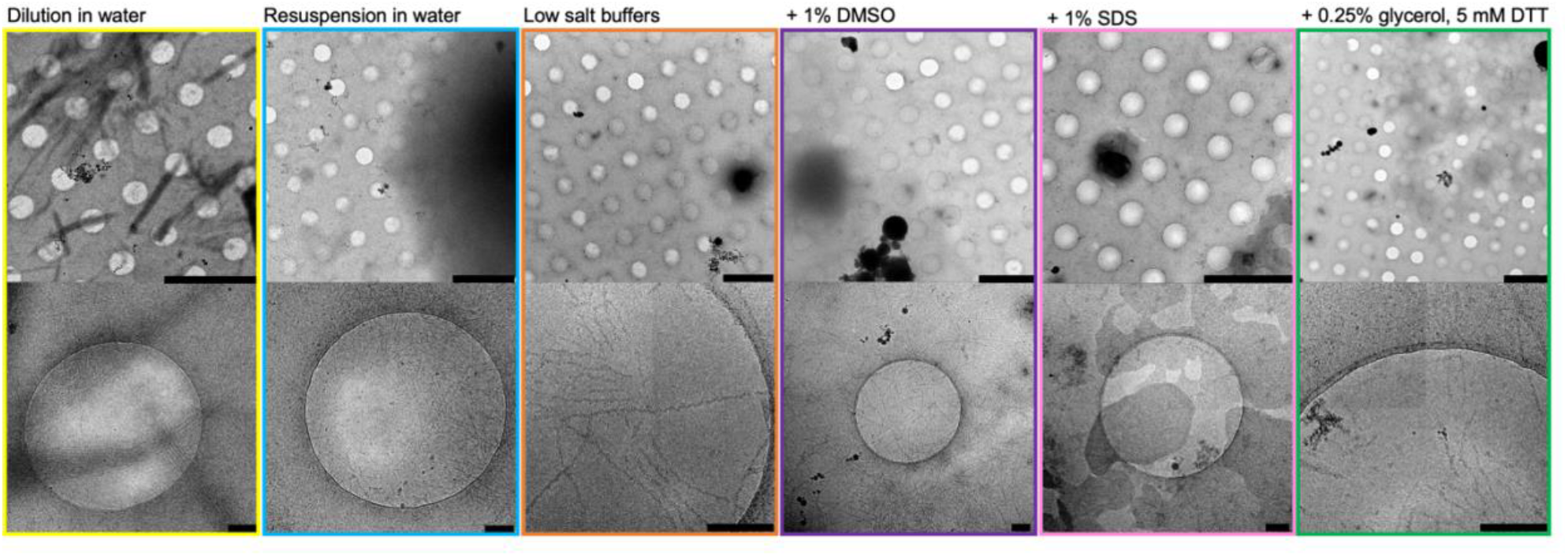
CryoEM grid optimization. Reversible fibrils were plunge frozen in various buffers for optimal particle dispersion and minimum interactions at the air-water interface (AWI), representative micrographs shown. Yellow and blue: dilution and resuspension in water caused significant clumping and dissociation. Orange: low salt buffers < 100 mM NaCl cause denaturation at the air water interface. Purple and pink: addition of DMSO and SDS, respectively, reduced clumping but impacted ice and fibril integrity. Green: final freezing condition 0.25% glycerol and 5 mM DTT resulted in acceptable clumping and reduced AWI denaturation. Scale bar top row = 2 μM, scale bar bottom row = 200 nm.

**Supplemental Figure 5.**
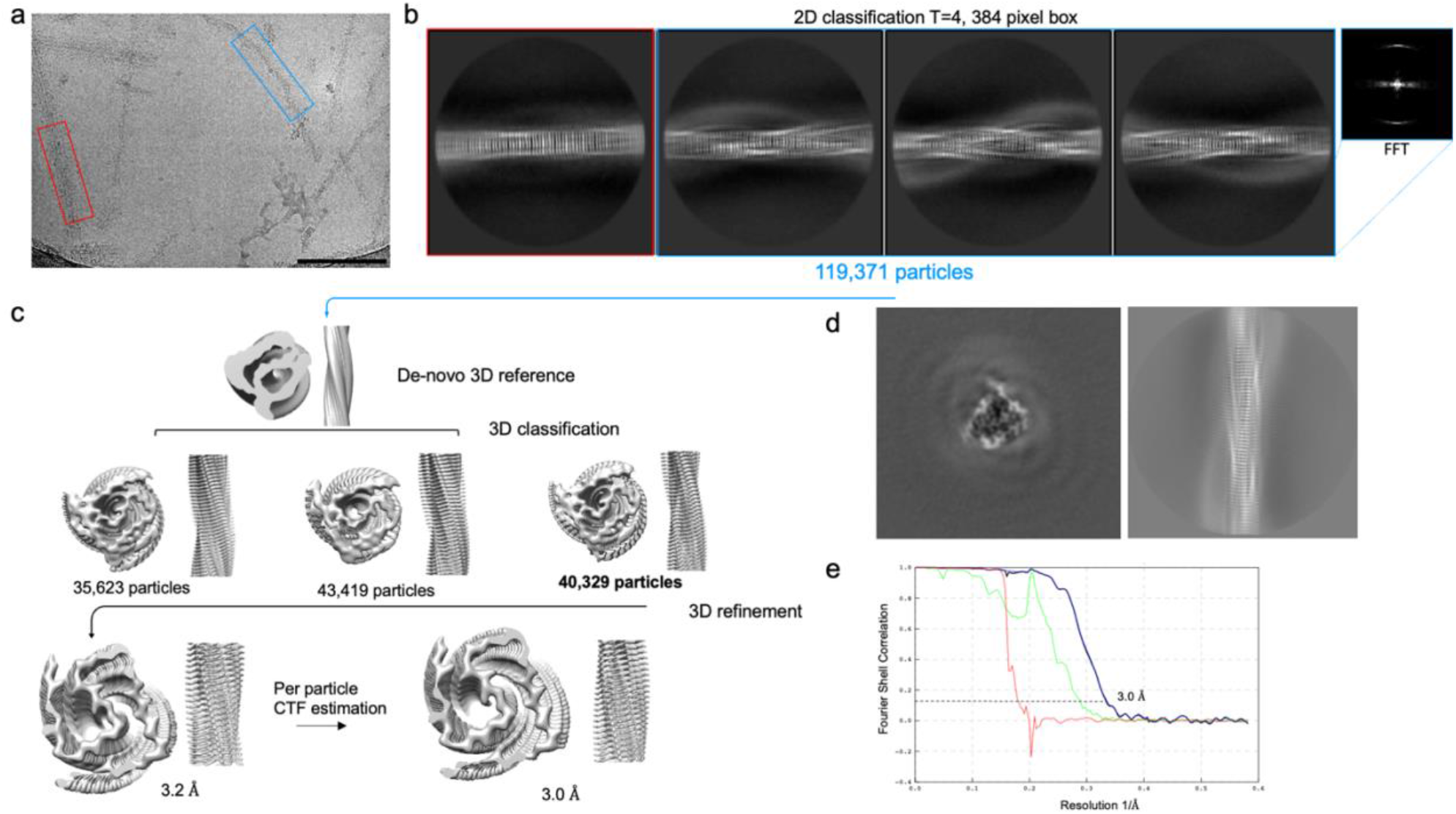
CryoEM data processing and model comparisons.**a**, CryoEM micrograph of twisting fibril polymorphs (blue) and non-twisting (red) Scale bar = 50 nm. **b**, Representative 2D class averages of polymorphs pictured in SF1a (left) and 4.8 Å reflection pattern in generated Fourier transform of 2D class (right). **c**, Schematic representation of helical reconstruction of major twisting species performed in RELION. **d**, Central slice from final reconstruction (left) and projection of 3D model (right). **e**, FSC curves between independently refined half-maps masked and corrected (black), unmasked and corrected (green), masked (navy blue) and phase randomized (red). Black dotted line indicates the FSC=0.143 resolution of 3.0 Å.

**Supplemental Figure 6.**
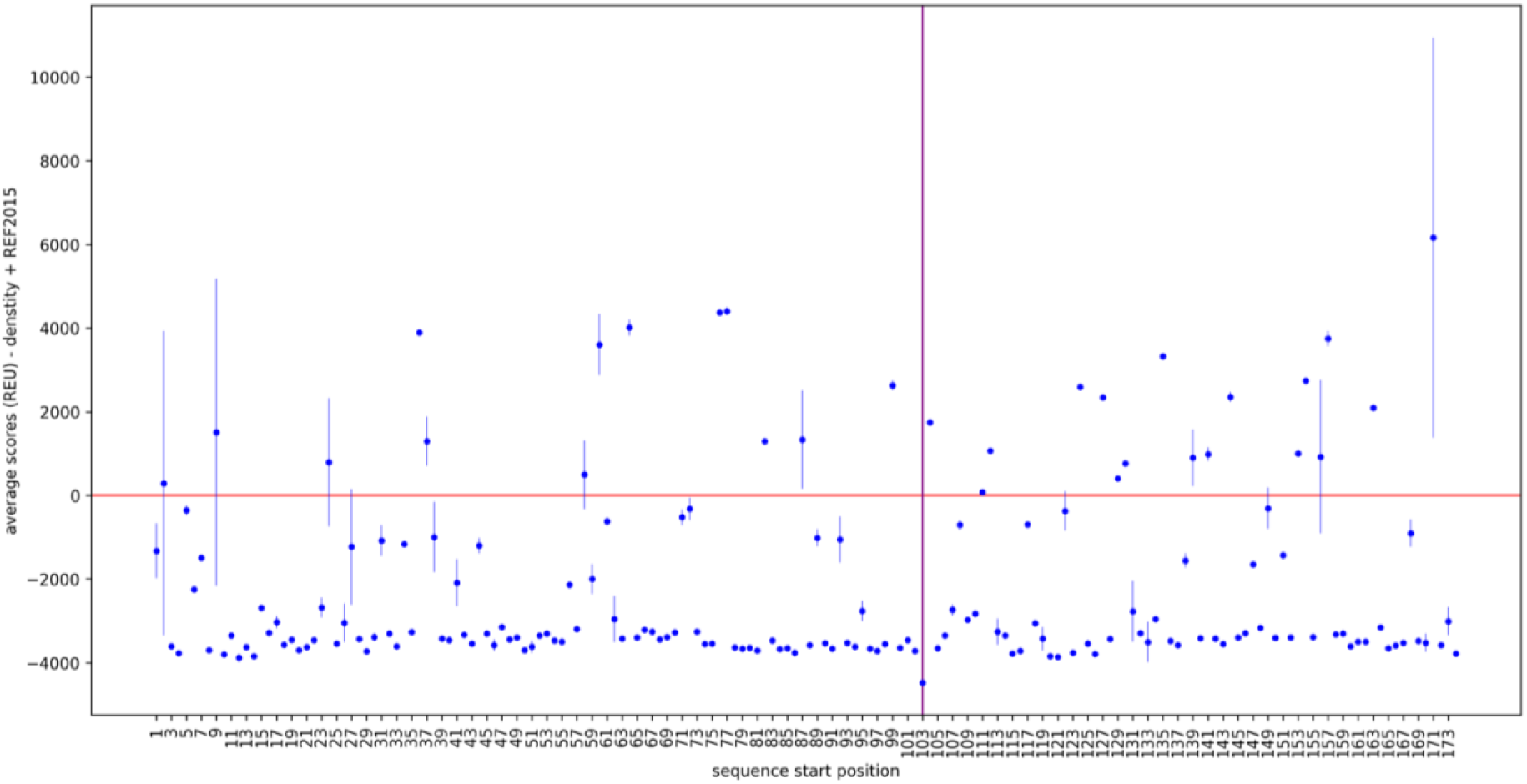
Average scores of threading experiment plotted against sequence start position. Average scores were calculated with Rosetta’s REF2015 energy function combined with a cryo-EM density score term. Each point represents the average of three trials. Sequence start position 103 shows the most favorable sequence window and is in agreement with the modeled sequence.

**Supplemental Figure 7.**
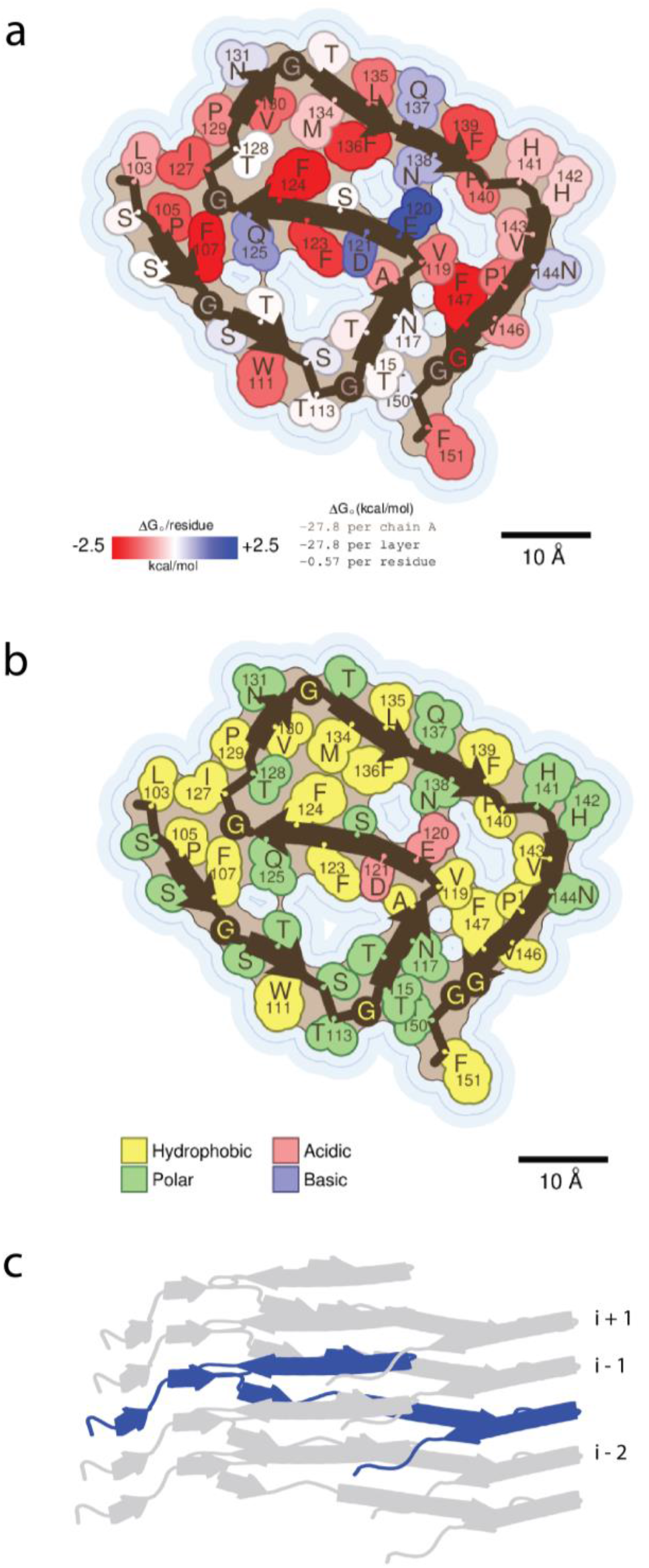
Structural features of CPEB3AC that may influence its stability. **a.**Energy per residue stabilization, key lower left. Thin blue lines represent the closest approach that the center of a water molecule can achieve. **b, c,** Cartoon rendering secondary structure elements and highlighting the height differences within a single chain that interact with 3 consecutive layers.

**Supplemental Figure 8.**
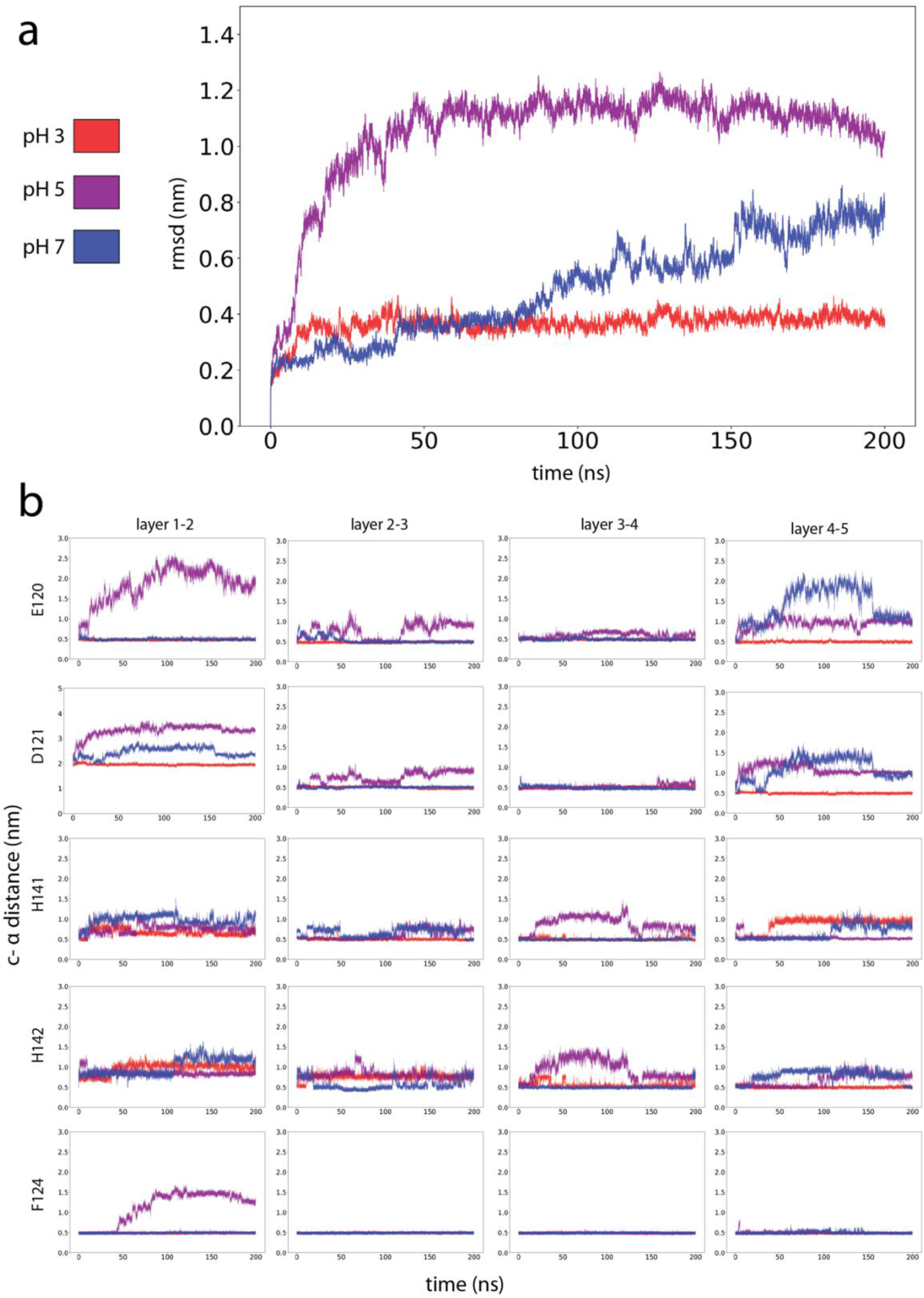
Molecular dynamics simulations of pH triggered instabilities in fibril core. pH color legend where pH 3 (red), pH 5 (purple) and pH 7 (blue).**a**, Averaged RMSD for all layers across 200 ns **b**, Selected charged residues from top to bottom: E120, D121, H141 and H142. Control residue F124. From left to right: neighboring layer c-α distances across 200 ns simulations.

**Supplemental Figure 9.**
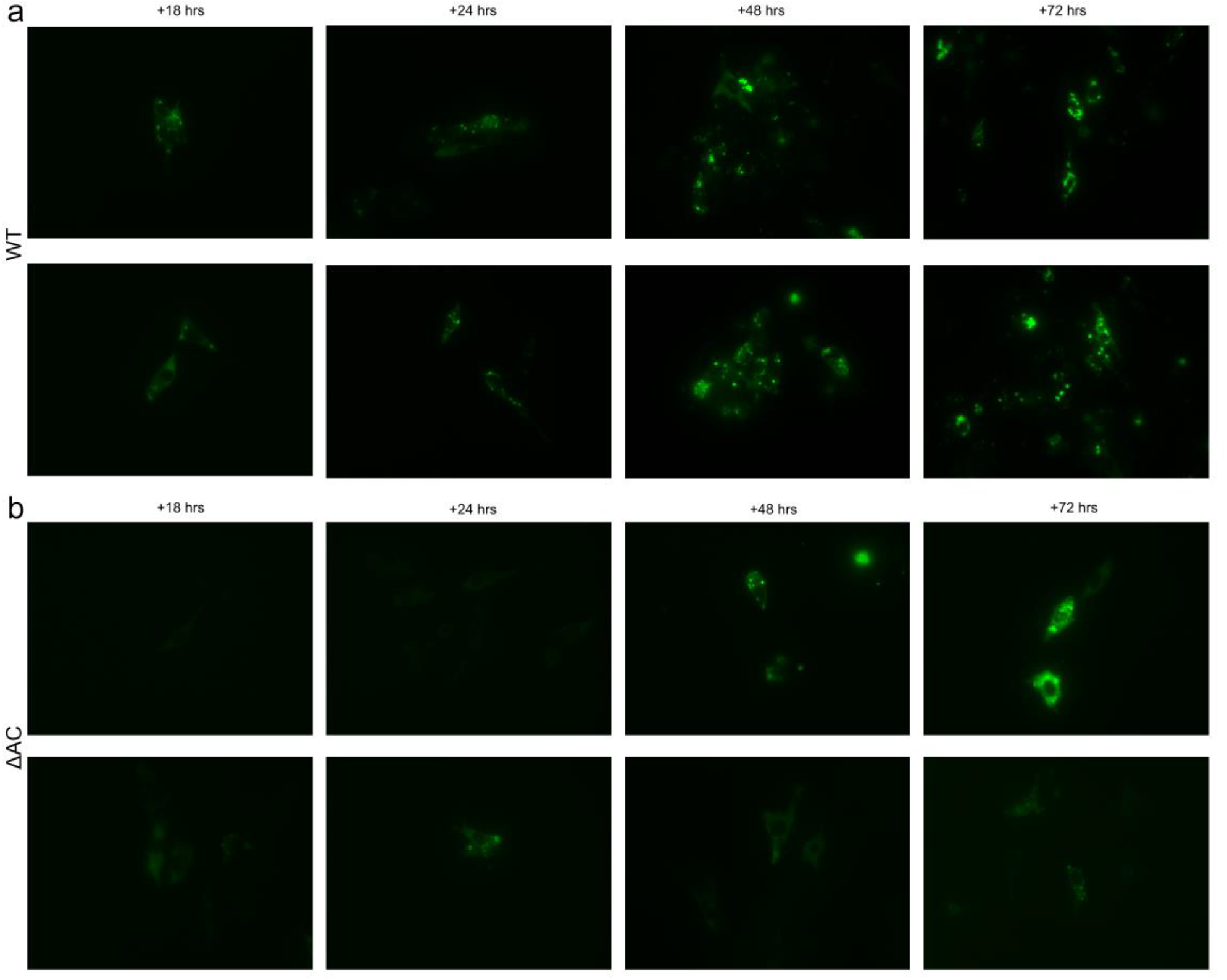
Aggregation phenotypes of CPEB3 WT and ΔAC constructs in HT22 in GFP channel (green).**a**, Representative images of WT CPEB3-GFP expression across four time points 18, 24, 48 and, 72 hrs. **b**, Representative images of CPEB3^ΔAC^-GFP expression across four time points 18, 24, 48 and, 72 hrs with earlier time points displaying significantly diffuse phenotypes. All time points indicate time after incubation with lipofectamine and corresponding vector.

**Supplemental Figure 10.**
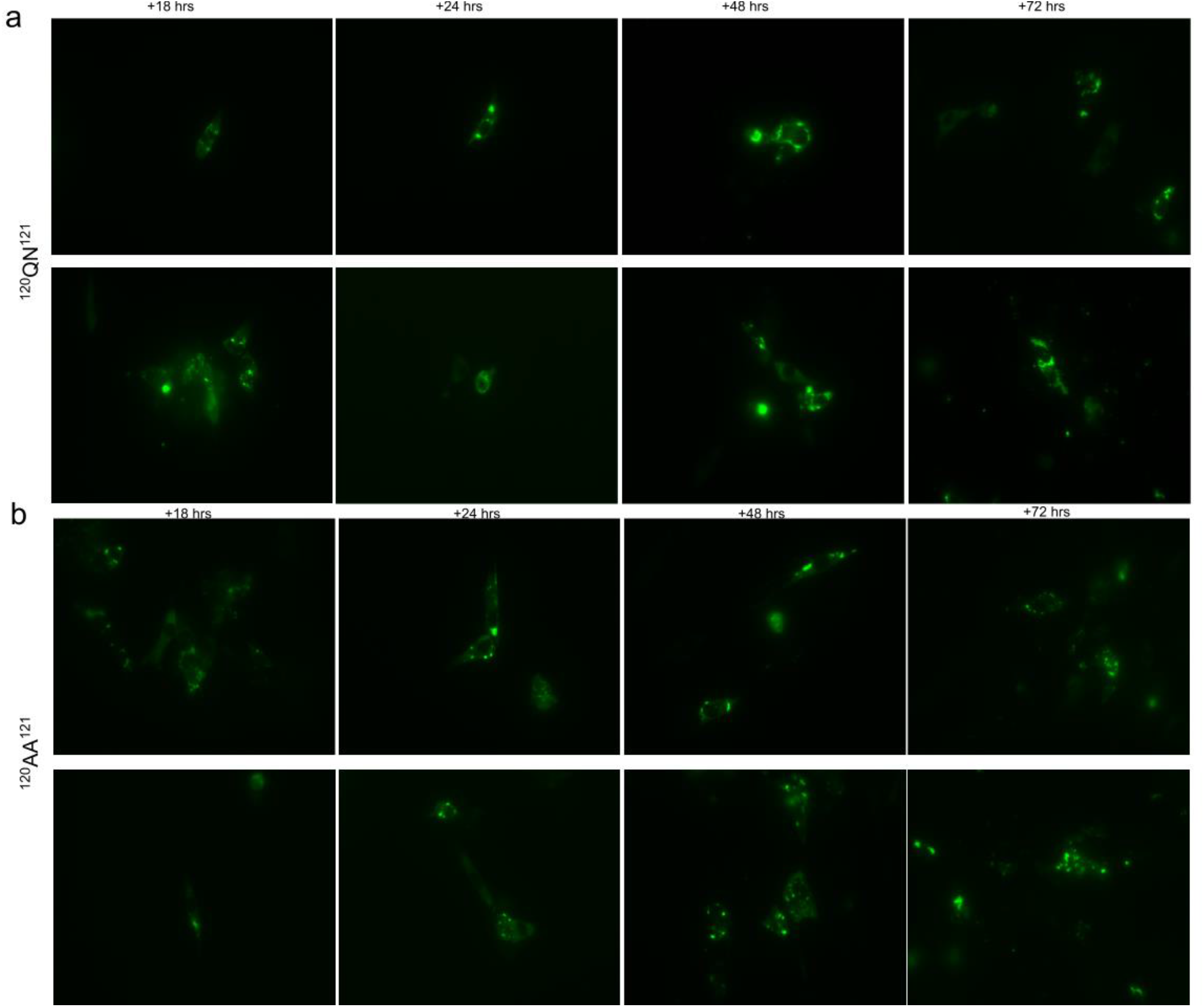
Irregular aggregation phenotypes of CPEB3 mutant constructs in HT22 in GFP channel (green).**a**, Representative images of CPEB3-^120QN121^-GFP across four time points 18, 24, 48 and, 72 hrs. **b**, Representative images of CPEB3^120AA121^-GFP expression across four time points 18, 24, 48 and, 72 hrs. As in Supplemental Figure 4, all time points indicate time after incubation with lipofectamine and corresponding vector.

**Supplemental Figure 11.**
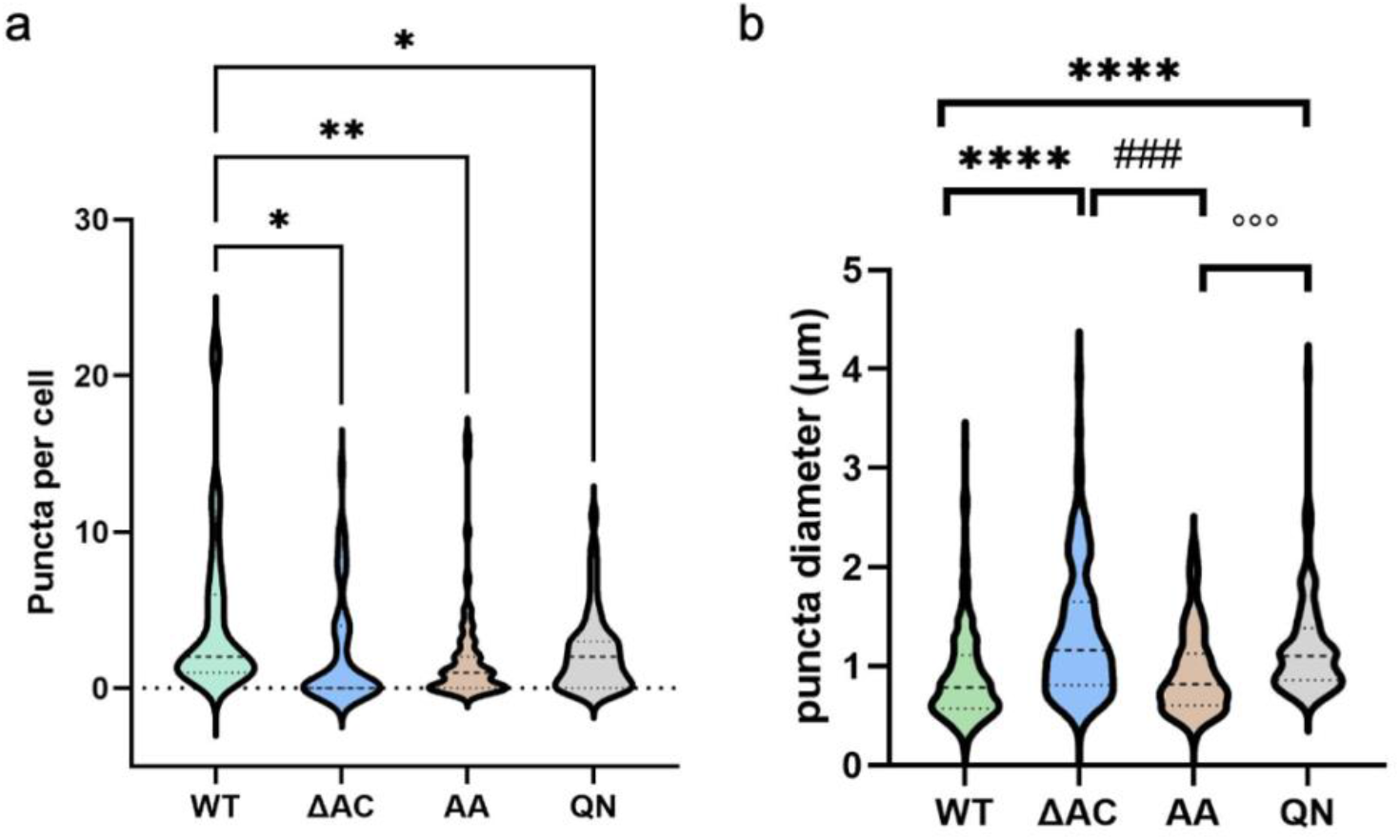
CPEB3 mutant phenotypes.**a**, Quantification of puncta per cell after 24 hours of exogenous expression. Where WT-GFP n=60 cells, CPEB3^ΔAC^-GFP n=52 cells, CPEB3-^120QN121^-GFP n=56 cells and CPEB3^120AA121^-GFP n=50 cells. One-way ANOVA, followed by Tukey’s multiple comparisons post hoc test, * p<0.05, ** p<0.01, vs WT. **b**, Quantification of average puncta diameter from cells surveyed in **a**. One-way ANOVA, followed by Tukey’s multiple comparisons post hoc test, **** p<0.0001, vs WT, ### p<0.001, vs dAC, °°° p<0.001 vs QN.

## Materials and Methods

### Construct design

Design of recombinant CPEB3 segment included previously reported residues critical for aggregate formation and prion-like behavior in cells^7^. CPEB3 residues 1-217 were inserted into

pET—24(+) vector from TWIST biosciences. Sequences were verified via DNA sequencing (Genewiz). CPEB3-EGFP constructs were a kind gift from Luana Fioriti.

**Panel 1:**
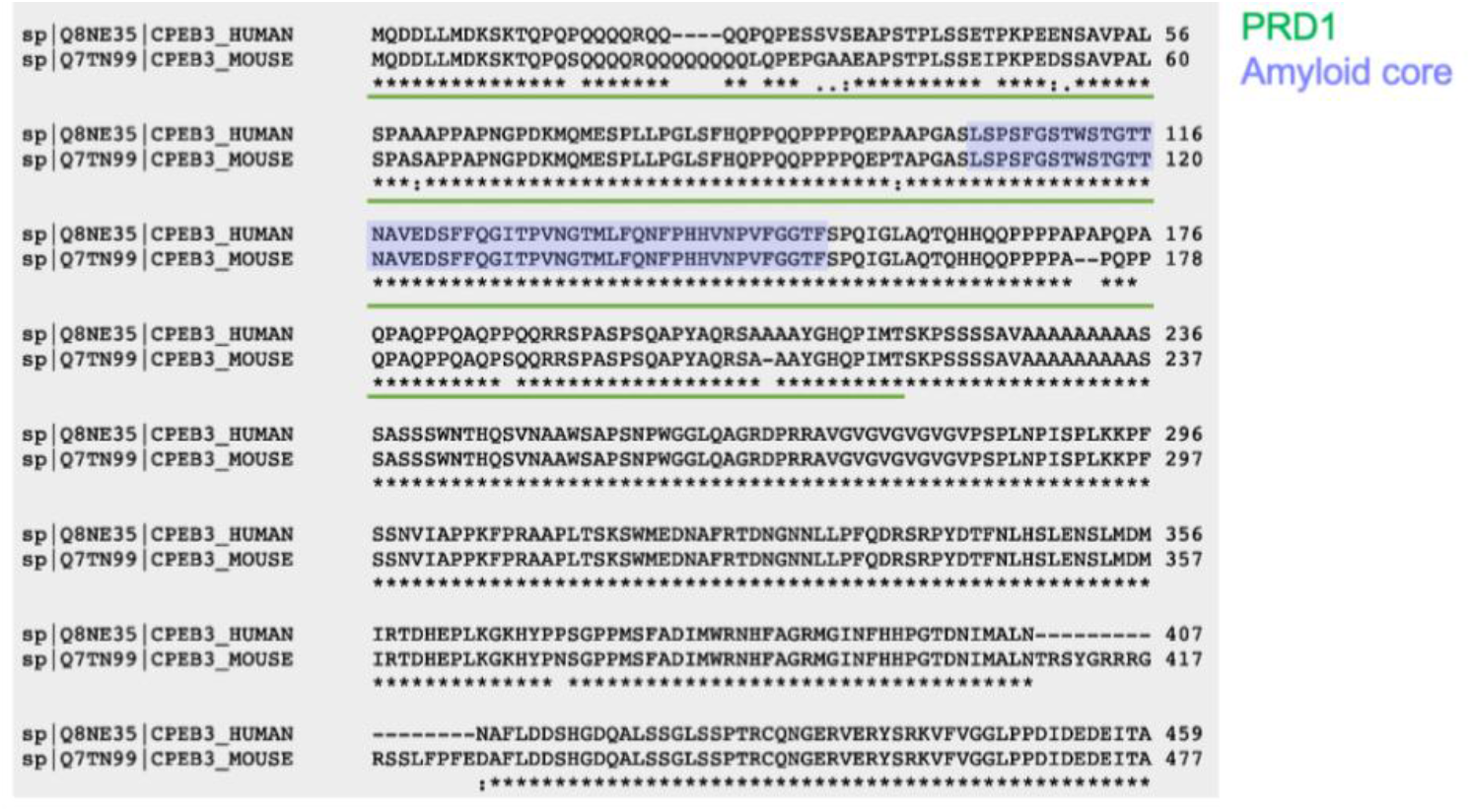
Sequence alignment of CPEB3 human and mouse sequences. Green light denotes residues within PRD1 and lilac highlight corresponds to residues placed in amyloid core structure.

### Protein expression and purification

Recombinant CPEB3^1-217^ was expressed in *Escherichia coli* BL21 GOLD cells. Briefly, cells were grown in LB media with 50 ug/ml kanamycin at 37° C to an OD600 of 0.6-0.8. 1 mM isopropyl B-D-1-thiogalactopyranoside (IPTG) was added for protein expression and cells were cultured an additional 6 hrs at 37° C. Cells were harvested and stored at -20° C until purification. Cell lysates were resuspended in lysis buffer containing 10 mM Tris, 1 mM EDTA at pH 8.2 and supplemented with 1% halt protease inhibitor cocktail (Thermo Scientific), 1% homemade Benzonase and subsequently lysed using 6 consecutive freeze-thaw cycles in liquid nitrogen. Cell lysates were centrifuged (12,000gs for 30min) and pellet was resuspended in 6M Gdn-HCl, 50mM Tris-Base, 25mM imidazole at pH 8 and left gently shaking on ice overnight. Lysates were centrifuged (3,000gs for 30 min) and supernatant was filtered and loaded onto an equilibrated HisTrap HP column (GE) using NGC (Biorad). The HisTrap was pre-equilibrated with 6M Gdn-HCl, 50mM Tris-Base and 25 mM imidazole. Protein was eluted with gradient protocol containing 6M Gdn-HCl, 50mM Tris-Base and 500 mM imidazole. Eluted protein was flash frozen and stored at -80° C until further purification. Further purification was executed with RP-HPLC, using an Interchim puriFlash® 4125 Preparative Liquid Chromatography System equipped with an Interchim PT-15C8-F0040 cartridge (C8,15 μm particle size). The eluted protein fractions were defrosted from -80°C and pooled. The pooled sample was then diluted to 1.8 mL with a dilution buffer consisting of 6 M Gdn-HCl, 50 μM NaH 2 PO 4, 10 μM Tris, and 20 μM imidazole. Before injection the 2-mL sample loop was washed with the above dilution buffer to maintain high concentrations of guanidine to minimize protein aggregation. [A] consisted of a 99 to 1 water to acetonitrile solution with 15 mM NH 4 OH added as a buffer. [B] consisted of a 40 to 60 water to acetonitrile solution with 15 mM NH 4 OH. Flow rate was set at 40 mL/min. Before injection, the system was equilibrated to 100% [A]. The protein sample was loaded onto the loop and injected. Three wavelengths were used to monitor the purification: 214nm (black), 280 nm (orange), and 254 nm (light blue). The gradient was held at 100% [A] for 5 minutes then a gradual increase to 100% [B] was executed over a span of 25 minutes (Δ5% per minute). The target protein started eluting at 16 minutes.

### Fibril preparation and negative stain TEM

Lyophilized protein powder was resuspended in H2O at 1.2 mg/mL concentration, aliquoted and immediately flash frozen and stored at -20°C until further use. Various buffers containing urea and guanidine were initially screened for fibril formation to attempt to control clumping and aggregation. These aggregates were heterogenous and difficult to reproduce. 96-well screen ranging in salt concentrations and pH were used for subsequent attempts without denaturants. The optimized fibril growth condition was incubation in 125mM NaCl, 50mM Tris-Base, 10mM K2PO4 and 5mM glutamic acid ∼pH 4.5 and left shaking overnight on the acoustic shaker. Fibrils were collected by scraping and pooling solution from wells and immediately prepared for subsequent experiments. Negative-stain transmission EM samples were prepped by applying 3 μl of fibril solution to glow-discharged 300 mesh carbon-coated formvar support films mounted on Cu grids (Ted Pella). The samples were wicked, washed briefly with H2O, stained with 2% uranyl acetate for 90 seconds and wicked again, allowing to air dry for 3 minutes. Grids were imaged on a T12 (FEI) electron microscope.

### Cryo-EM sample preparation and data collection

Freshly harvested fibril solution was diluted 5 fold with buffer containing 100mM NaCl, 10mM Tris-Base, 5mM DTT and 0.25% glycerol. 1.6 μL of diluted fibril solution was applied to each side of a glow-discharged Ultrathin carbon on Quantifoil 1.2/1.3 300 mesh on Au grid (Ted Pella) and plunge frozen into liquid ethane using a Vitrobot Mark IV (FEI) set at 100% humidity, 4°C with a blot force of -1 and blot time of 1.5 seconds. Data were collected on a Titan Krios G3i microscope with a K3 direct detection camera (Gatan) operated at an accelerating voltage of 300 keV and a 20eV slit width (BioQuantum). Automated data acquisition were performed using EPU 2.8 software (Thermo Scientific). Images were acquired with a nominal magnified pixel size of 0.86 Å/pixel, a total of 40 frames with a dose of 1.3 electrons per Å^2^ per frame resulting in a total dose of 52 electrons per Å^2^.

### Data processing and helical reconstruction

CTF estimation were performed with CTFFIND 4.1.8. Drift-correction and dose-weighting were performed using MotionCorr2 implemented in RELION 3.1^36^. Particles were picked with the automatic particle picking software crYOLO^37^. All subsequent image processing and helical reconstruction were performed in RELION as previously described.^29,38^ Briefly, fibril segments were initially extracted with a 90% overlap into 512 pixel boxes binned by 2 (1.72 A/pix) and subjected to reference-free 2D classification using a T=2 regularization parameter. Segments belonging to suboptimal 2D averages were discarded and homogenous subsets were selected for further processing. Upon identification of a rapidly twisting species, a smaller box size was chosen. All segments were then extracted with a 90% overlap into 384 pixel boxes for a total of 2,410,032 segments. Segments were split into 10 groups and several rounds of reference free 2D classification were performed to weed out suboptimal segments. Homogenous subsets from each were regrouped and reference free 2D classification were iteratively run decreasing psi_step and offset_step from 8 to 1. After the majority of suboptimal particles had been discarded an additional round of classification were performed with the tau regularization parameter set to T= 8. Only segments contributing to averages showing clear 4.8 Å signal in corresponding 2D FFTs were selected for subsequent 3D processing, resulting in 119,372 segments. Using both the 512 pixel box and 384 pixel box an initial helical pitch of 536 Å and helical twist of -3.22° were estimated. Starting with these estimates a 3D reference was reconstructed de-novo with 384 pixel 2D averages showing clear β-strand separation along the helical axis. 3D manual refinements were performed using a 30 Å low-pass filtered initial model, K=3 and with manual control of tau_fudge and healpix to reach a resolution of ∼ 5Å. Segments contributing to a homogeneous class as defined by stable helicity and separation of β-strands resulted in a final subset of 40,329 segments. Again, a manual 3D refinement was performed on selected segments leading to a reconstruction of 3.5 Å. Several 3D auto-refinements using a 7Å low-pass filtered map from the final manual refinement were run with optimization of helical parameters leading to an estimated helical rise of 4.91 Å and a helical twist of -3.32°. This map was then used for per-particle CTF estimation and a 3D auto-refinement was repeated. The nominal pixel size was adjusted from 0.86 Å to 0.879 Å, resulting in an expected helical rise of 4.8 Å. Post-processing of the map with an extended initial mask of 3 pixels and soft edge of 10 pixels was followed by clipping and centering the map using clip.com. Auto-sharpening using phenix.auto_sharpen at the resolution cutoff indicated by the half-map GSFSC led to a final overall resolution of 3.0 Å. The atomic model was built into the refined map using COOT^39^. We performed automated structure refinement using phenix.real_space_refine (Phenix 1.2)^40^.

### Threading

Threading was performed with a custom python script utilizing the PyRosetta^41^ software package and a sliding window approach. For each window, the experimentally expressed sequence was threaded onto a poly-alanine backbone then energy minimized with the PyRosetta FastRelax with cryo-EM density function. FastRelax used the Ref2015^42^ energy function with a modified fa_elec weight to 1.5 and elec_dens_fast weight of 25. Each sequence window was energy minimized in triplicate and the resulting scores were averaged. Symmetry was applied to pose objects to increase speed of computation.

### Energetic calculations

Solvation free energy calculations were performed as recently described^18^. Briefly, the solvent accessible surface area (SASA) for each atom in a residue was determined (folded state). Next, the SASA for each atom of a residue was determined in the absence of the other residues (reference state), and the difference between both states was calculated (SASA_Ref_ – SASA_Fold_). This value was then multiplied by the Atomic Solvation Parameter (ASP) specific to each atom, as determined previously by Eisenberg et al^43^. An entropic term is also included to take into consideration the degrees of freedom lost in going from a disordered to ordered state^44^. The energies of all atoms were then summed to generate the solvation energy for each structure. Difference energy maps were generated by subtracting solvation free energies pairwise for each atom in the two structures being compared.

### Molecular dynamics simulations

Molecular dynamics simulations were performed with the GROMACS^45^ software package (version 2022.2) using the CHARMM27^46^ all-atom forcefield. Simulations were carried out with a 5-layer CPEB fibril under varying charge states to simulate fibril dynamics at pHs 3, 5, and 7. In each case, the fibril was placed in a cubic box, solvated with SPC/E three-point water molecules and added counter ions. The system was energy minimized, then temperature and pressure equilibrated for 100 ps. Final simulations were executed for 200 ns.

### Thioflavin T-binding and dissolution assays

Protein was resuspended in fibril formation buffer as described previous at 25 μM concentration with equal parts ThT. Solution was pipetted into a 96-well well plate with optical bottom (Sigma-Aldrich) and incubated at 37°C with continuous shaking. ThT fluorescence was measure with an excitation filter of 440 nm and emission filter of 480nm using a Varioskan plate reader (ThermoFischer). Aggregation curves were generated from triplicate, n=3 wells. Dissolution assays were started after 18 hours of fibril formation. Because fibrils were not able to be resuspended in different buffers due to their liability, a 96-well plate was prepared by diluting each fibril condition 3-fold in corresponding buffers. Normalization of ThT intensity was performed to account for variations across experiments and reduction of signal when diluting samples for dissolution assays.

### Cell culture

HT22 hippocampal neurons and U87 glioblastomas were cultured in Dulbecco’s modified Eagle’s medium with 10% Fetal Bovine Serum (ThermoFisher) and 1% penicillin/streptomycin (ThermoFisher) in a tissue-culture incubator at 37°C and 5% CO2. Cells were cultured as described above and plated on 12-mm glass coverslips coated with poly–L-lysine (Sigma) and Lamin 50 uG (Sigma) in 6-well cell-culture plates. Cells were plated at 70% confluency overnight and transfected using Lipofectamine 3000 kit (ThermoFisher) for an 8 hr incubation. Cells were harvested and/or imaged at 24 hours post transfection for all experiments. Live fluorescent images were acquired on an EVOS M7000 (ThermoFisher) on a 40X objective.

### Immunofluorescence and puncta analysis

Cells were fixed in 2% paraformaldehyde at 4°C for 10 minutes and permeabilized using 0.5% PBS-Triton X-100 for 10 minutes. Cells were subsequently washed with PBS-T and blocked with 2% BSA, 0.5%FSG, and PBS-T solution for 1hr at RT. Cells were incubated with 1° antibodies diluted in blocking solution for approximately 2 hours. Cells were washed with PBS-T three times before beginning incubation with 2° antibodies and washed with PBS-T. Coverslips were mounted using Prolong Glass Antifade Mountant (ThermoFisher) and allowed to dry for 72 hours. Fluorescent images were acquired on a Leica confocal SP8-STED/FLIM/FCS on a 60X water immersion objective. Puncta were manually counted and distances were manually calculated by averaging X and Y for each puncta in GFP channels. Image analyses were performed in FIJI (ImageJ).

### Luciferase assays

HEK293 cells were plated at 50% confluency overnight and transfected using Lipofectamine 3000 kit (ThermoFisher) with ∼0.5 μg of CPEB3 DNA, 0.5 μg of Renilla luciferase appended with GluA2 3′ UTR or SUMO2 3’UTR, and 0.5 μg of Firefly luciferase^47^. 18/24 hr after transfection, cells were lysed with 100 μl of buffer for dual luciferase assay (Promega). 20 μl of the lysates were used for the quantification of Firefly and Renilla Luciferase activity using a Luminometer (GloMax, Promega).

Hippocampal neurons cultured for 9–10 days in Neurobasal medium with B27 supplement (Invitrogen) at a cell density of 30 000–40 000/cm2 were cotransfected (Lipofectamine LTX, LifeScience) for 3 h with ∼0.5 μg of CPEB3 DNA, 0.5 μg of Renilla luciferase appended with GluA2 3′ UTR, and 0.5 μg of Firefly luciferase. We used glycine to stimulate the NMDA receptors selectively, a protocol that mimics LTP induction in cultured neurons. Chem-LTP was induced as described previously^48^. Briefly, neuronal cultures were transferred from Neurobasalgrowth medium to extracellular solution (ECS) containing: 150 mM NaCl, 2 mM CaCl 2, 5 mM KCl, 10 mM HEPES(pH 7.4), 30 mM glucose, 0.5mM TTX, 20mM bicuculline methiodide. The transfected neurons were stimulated with 200μM glycine for 3 min and then washed for 30 minutes before lysis in 100 μl of buffer for dual luciferase assay (Promega). 20 μl of the lysates were used for the quantification of Firefly and Renilla Luciferase activity using a Luminometer (GloMax, Promega). The ratio between Renilla and Firefly is calculated for each experimental condition.

### Statistical analyses

Number of repeats and (Avg) ± SE are stated in figure legends. Statistical significance was measured by one or two-way ANOVA followed by the Tukey’s multiple comparisons test.

## Acknowledgments

We are grateful to Wong Hoi Hui of the Electron Imaging Center for NanoMachines (EICN) for help and advice during cryoEM screening and sample preparation. We thank Peng Ge (UCLA) for helpful discussions during cryoEM data processing, assistance during data acquisition and Duilio Cascio (UCLA) for assistance during model building and refinement.

We thank Corey Hecksel (SLAC) for assistance during data acquisition. Part of this work was performed at the Stanford-SLAC Cryo-EM Center (S2C2), which is supported by the National Institutes of Health Common Fund Transformative High-Resolution Cryo-Electron Microscopy program (U24GM1295410). M.D.F is supported by Eugene V. Cota-Robles Fellowship, Ruth L. Kirschstein NRSA GM007185, Whitcome Pre-Doctoral Fellowship, Audree V. Fowler Fellowship and a National Science Foundation Graduate Research Fellowship. J.A.R. is supported as a Pew Scholar, a Beckman Young Investigator and a Packard Fellow. This work was performed as part of STROBE, an NSF Science and Technology Center through Grant DMR-1548924, and by DOE Grant DE-FC02-02ER63421. The work was also supported by NIH-NIGMS Grant R35 GM128867.

## Author Contributions

M.D.F designed experiments, purified recombinant constructs, screened negative stain samples, performed ThT assays, prepared cryoEM samples, performed cryoEM data collection and processing helical reconstruction, purified vectors for mammalian expression, cultured mammalian cells, performed live fluorescent imaging, immunofluorescent experiments and performed data analysis. M.R.S performed solvation energy calculations and assisted in model building and refinement. S.Z. performed computational threading and MD simulations. D.R.B assisted in cryoEM data processing helical reconstruction. S.T and A.S performed puncta quantification. T.Z. and J.C. assisted in recombinant protein expression and purification. L.F. cultured primary neurons, performed luciferase assays and analysis and guided the project. M.D.F and J.A.R wrote the manuscript with input from all authors. J.A.R supervised and guided the project.

